# Metabolic Responses of Different Levels of Fitness

**DOI:** 10.64898/2025.12.27.696707

**Authors:** Inigo San-Millan, Janel L. Martinez

## Abstract

The metabolic and physiological responses to exercise vary markedly across different levels of fitness and training status. While maximal oxygen uptake (VO₂max) has historically been used as the primary indicator of cardiorespiratory fitness, increasing attention has been directed toward metabolic and bioenergetic responses to exercise, including blood lactate concentration and substrate utilization.

In this study, we examined comprehensive physiological and metabolic responses during graded exercise testing in 204 male cyclists spanning four fitness categories: Tour de France professional cyclists, competitive cyclists, master cyclists, and recreational cyclists.

Measurements included power output, VO₂, blood lactate concentration, and rates of fat and carbohydrate oxidation derived from indirect calorimetry.

Across all exercise intensities, performance and metabolic parameters followed a clear hierarchical pattern corresponding to competitive level. Tour de France cyclists demonstrated significantly greater power output, higher VO₂max, lower blood lactate concentrations at matched workloads, higher fat oxidation rates, and delayed reliance on carbohydrate oxidation compared with less-trained groups. Strong inverse correlations were observed between blood lactate concentration and fat oxidation, while positive correlations were observed between blood lactate concentration and carbohydrate oxidation across all fitness levels.

## 1. INTRODUCTION

The metabolic response to exercise has been a major area of interest among physiologists since the end of XVII century when the father of modern chemistry, Antoine Lavoisier, studied for the first time the amount of oxygen (O2) consumed during exercise(Underwood, 1944). During the first part of XX century, a high level of interest in understanding the physiological and metabolic responses to exercise emerged. Early investigations focused primarily on oxygen consumption, substrate utilization, and lactate production. Fletcher and Hopkins showed in 1907 that contracting frog muscles produced lactate which disappeared upon exposure to oxygen-rich environment(Fletcher, 1907). This phenomenon was attributed to O2 exposure and named “O2 Debt” in 1923 by A.V. Hill and Lupton(Hill, 1923). Then, the pioneer work in the 1930’s by Francis Gano Benedict regarding fat and carbohydrate metabolism during exercise, resulted in novel understanding in the metabolic responses to exercise(Benedict, 1938).

In the last three decades, metabolic testing has become increasingly popular among competitive athletes to assess fitness levels and prescribe individualized training programs aimed at improving performance. Historically, the assessment of maximal oxygen consumption (VO₂max) has been considered the “gold standard” for evaluating cardiorespiratory responses to exercise(Shephard, 1968). However, as knowledge of exercise metabolism has advanced, the interest of researchers and applied physiologists has expanded toward the study of metabolic and bioenergetic responses to exercise, beyond cardiorespiratory capacity alone.

One of the main effects of endurance training at the cellular level is increased mitochondrial content and oxidative capacity, accompanied by enhanced lactate clearance(Dubouchaud et al., 2000, McDermott and Bonen, 1993, Donovan and Brooks, 1983, Bergman et al., 1999, San-Millan and Brooks, 2018, Bonen et al., 1998, Gollnick et al., 1986). Consequently, measuring blood lactate concentration ([La⁻]) has been widely adopted by physiologists and coaches to evaluate metabolic responses to exercise, monitor training adaptations, and prescribe individualized training programs(Jacobs et al., 1983, Billat, 1996).

The evaluation of substrate utilization (fats and carbohydrates) during exercise using stoichiometric equations via indirect calorimetry (Frayn, 1983) has also gained popularity in recent decades for assessing metabolic responses during exercise (San-Millan and Brooks, 2018, Randell et al., 2017, Achten and Jeukendrup, 2003, Achten et al., 2003, Stisen et al., 2006)(Frandsen et al., 2017, Knechtle et al., 2004). Furthermore, the combination of measuring [La^−^] and fat oxidation rates (FATox) can be a useful method to indirectly assess mitochondrial function in athletes as we have recently proposed(San-Millan and Brooks, 2018).

The sport of cycling has been at the forefront of applied exercise physiology and scientific approaches to training. Numerous studies have examined physiological responses in competitive cyclists, including measurements of blood lactate concentration, VO₂max, and power output (Lucia et al., 2000a, Lucia et al., 2000b, Lucia et al., 1998, Hoogeveen, 2000, Impellizzeri et al., 2008, O’Toole et al., 1989, Padilla et al., 1999, Padilla et al., 2001, San-Millan et al., 2009, San-Millan and Brooks, 2018). These studies have measured different parameters including [La^−^], VO2_max_, or power output, alone or in conjunction, to assess performance. However, many of these studies have involved relatively small cohorts or single teams, and comprehensive metabolic assessments incorporating substrate utilization have been less commonly reported despite their relevance for performance evaluation.

In the present study, we report comprehensive physiological and metabolic data obtained during graded exercise testing in cyclists spanning a broad range of fitness levels, from Tour de France professionals to recreational cyclists. Parameters assessed included VO₂, blood lactate concentration, substrate utilization, and power output.

Additionally, we examine the relationships between blood lactate concentration and rates of fat and carbohydrate oxidation across exercise intensities, highlighting the practical utility of lactate testing for characterizing metabolic responses to exercise and informing performance assessment.

## 2. METHODS

### Materials and Methods

#### Description of subjects

204 male cyclists, separated into four cohorts, performed graded exercise tests to volitional exhaustion on a leg cycle ergometer (Lode Excalibur, Lode, The Netherlands). The cohorts were comprised of Tour de France professional cyclists (TdF), n = 23; competitive cyclists (CC), comprised of domestic professional cyclists and elite amateurs, n = 51; master cyclists (MC) >50 years old, n = 29; and recreational cyclists (RC), n = 101 (Table 1). The requirement to qualify for the TdF cohort was to compete at the highest level of professional road cycling, which is the Tour de France. To be eligible for the CC group, cyclists must compete at a national professional or elite-amateur level. The requirement for the MC cohort was to compete in the category of “masters” in which cyclists must be over fifty years old. Finally, the requirements for the RC group was to exercise at least 3 times/week and a minimum of 150 min per week for at least half a year. All subjects were given nutritional recommendations to consume >50% of total caloric intake as carbohydrates the evening before and the day of testing. Subjects were advised to avoid intense exercise or exercise exceeding two hours the day before testing and to refrain from exercise on the day of the test. This protocol was designed to reflect typical applied physiology testing conditions rather than strict overnight fasting.

**Table 1.** Characteristics of study cohorts.

| Parameter | Recreational<br>(N = 111) | Competitive Masters<br>(N = 30) | Competitive<br>(N = 51) | Tour de France<br>(N= 23) |
| --- | --- | --- | --- | --- |
| **Age (years)** |  |  |  |  |
| N | 101 | 29 | 51 | 23 |
| Mean ± SD | 45.8 ± 13.1 | 46.6 ± 6.1 | 29.8 ± 7.8 | 26.9 ± 2.7 |
| Min – Max | 18 – 72 | 40 – 60 | 18 - 39 | 21 – 32 |
| **Height (cm)** |  |  |  |  |
| N | 101 | 29 | 51 | 5 |
| Mean ± SD | 177.7 ± 6.9 | 178.3 ± 4.9 | 181.2 ± 6.0 | 181.6 ± 4.2 |
| Min – Max | 151.1 – 194.3 | 165.1 – 188.0 | 167.6 – 193.0 | 176.5 – 188.0 |
| **Weight (kg)** |  |  |  |  |
| N | 111 | 30 | 51 | 23 |
| Mean ± SD | 76.1 ± 8.4 | 72.4 ± 6.0 | 71.7 ± 7.1 | 69.2 ± 5.2 |
| Min – Max | 51.8 – 103.0 | 61.8 – 86.4 | 60.9 – 88.1 | 61 .0 – 76.1 |
| **Body Fat (%)** |  |  |  |  |
| N | 97 | 28 | 48 | 23 |
| Mean ± SD | 15.5 ± 4.1 | 13.2 ± 2.6 | 11.0 ± 2.2 | 9.7 ± 0.7 |
| Min – Max | 10.0 – 30.0 | 10.7 – 17.7 | 8.4 – 16.5 | 8.0 – 11.0 |

#### Hierarchy description

Throughout the results and discussion sections we employed the term “hierarchy” in order to classify each cohort according to the level of both performance and competitive level. Tour de France cyclists have reached the highest level possible in cycling and are at the top of the “hierarchy” followed by competitive cyclists, competitive master cyclists, then recreational cyclists.

#### Cycling Protocol

Each cyclist reported to the laboratory having abstained from food for at least two hours prior to testing and refrained from alcohol and caffeine for ≥8 hours. None of the cyclists followed a carbohydrate-restricted diet. All cyclists abstained from strenuous physical activity for 24 hours prior to testing. After a standardized 15-minute warm-up (<100 W), participants began a graded leg-cycling protocol. The protocol began at an intensity of 1.5 W·kg⁻¹ with increments of 0.5 W·kg⁻¹ every 10 minutes until volitional exhaustion, as previously described(San-Millan et al., 2009, San-Millan and Brooks, 2018). For recreational cyclists, the protocol began at 1.0 W·kg⁻¹, while for TdF cyclists it began at 2.0 W·kg⁻¹ for practical reasons. The goal was to observe the metabolic responses to exercise at a wide range of exercise intensities. Heart rate was monitored during the entire test with a heart monitor (Polar S725x, Polar Electro, Kempele, Finland). Study procedures were conducted in accordance with the Declaration of Helsinki and in accordance with a protocol that was approved by the Colorado Multiple Institutional Review Board (COMIRB)

### 2.1 Gas exchange measurements

Oxygen consumption (VO_2_), carbon dioxide production (VCO_2_), and Respiratory Exchanged Ratio (RER=VCO_2_/VO_2_) were determined using a ParvoMedics TrueOne 2400 Metabolic Measurement System (ParvoMedics, Inc.; Sandy, UT, USA). Subjects were required to wear a specialized mouthpiece that collected respiratory gases; respiratory gas data was averaged over 15 seconds throughout the entire test.

#### Fat and carbohydrate oxidation rates measurement

For the measurement of total body fat (FATox) and carbohydrate (CHOox), stoichiometric equations were applied to gas exchange measurements according to the methodology described by Frayn(Frayn, 1983):

CHO oxidation (g/min) = 4.55 × VCO_2_ – 3.21 × VO_2_
Fat oxidation (g/min) = 1.67 × VO_2_ – 1.67 × VCO_2_

### 2.2 Blood Lactate concentration measurement

At the end of every stage for the duration of the test, a sample of capillary blood was collected from the earlobe to analyze both intra- and extra-cellular levels of L-lactate (YSI 1500 Sport, YSI, Yellow Springs, OH, USA). Heart rate was monitored during the entire test with a heart monitor (Polar S725x, Polar Electro, Kempele, Finland).

### 2.3 Statistical Analysis

Data were analyzed using IBM SPSS Statistics version 26 (IBM, Armonk, NY, USA).

One-way ANOVA was performed to assess differences among groups. Levene’s test was used to assess homogeneity of variance. When assumptions were violated, Welch’s correction was applied. Bonferroni post hoc tests were used for multiple comparisons. This analysis was applied to each common parameter/timepoint within continuous data, represented graphically, reflecting the nature of the graded exercise assessment. Bivariate correlations to assess the statistical significance of relationships among variables studied over time were determined via Pearson’s correlation coefficient (r). Statistical significance was set at p < 0.05(*). Additionally, p < 0.01(**) and p < 0.001(***) are also reported. Data was graphed using Prism version 8 (GraphPad Software, San Diego, CA, USA).

## 3. RESULTS

As expected, Tour de France cyclists demonstrated superior performance compared with all other groups across measured parameters, with values generally following the hierarchical pattern TdF > CC ≥ MC > RC. Both absolute and relative power output were significantly better in TdF group compared to the rest of the groups (p<0.001, Table 2; Fig-1A-B).

**Table 2.** Parameter averages ± SD at each exercise intensity/stage for each fitness category. Parameters include power output (absolute Watts), blood lactate concentration (mmol/L), and VO_2_ (ml/kg/min). Body fat % according to Faulkner (Faulkner, 1966).

| Stage | Recreational<br>(N = 111) |  |  |  | Competitive Masters<br>(N = 30) |  |  |  | Competitive<br>(N = 51) |  |  |  | Tour de France<br>(N = 23) |  |  |  |
| --- | --- | --- | --- | --- | --- | --- | --- | --- | --- | --- | --- | --- | --- | --- | --- | --- |
| W/kg | N | [Watts] | [Lac.] | VO2 | N | [Watts] | [Lac.] | VO2 | N | [Watts] | [Lac.] | VO2 | N | [Watts] | [Lac.] | VO2 |
| 1.00 | 35 | 81.1<br>± 11.3 | 1.4<br>± 0.6 | 19.3<br>± 2.7 |  |  |  |  |  |  |  |  |  |  |  |  |
| 1.50 | 97 | 115.4<br>± 15.1 | 1.5<br>± 0.6 | 24.1<br>± 2.7 | 16 | 107.4<br>± 8.2 | 1.3<br>± 0.4 | 23.5<br>± 3.2 | 13 | 114.9<br>± 11.5 | 1.1<br>± 0.3 | 24.1<br>± 3.0 |  |  |  |  |
| 2.00 | 108 | 150.1<br>± 18.6 | 2.0<br>± 1.1 | 29.6<br>± 3.0 | 29 | 141.8<br>± 10.6 | 1.4<br>± 0.5 | 29.4<br>± 3.1 | 48 | 140.2<br>± 13.1 | 1.1<br>± 0.3 | 29.5<br>± 2.2 | 22 | 138.1<br>± 10.6 | 0.7<br>± 0.1 | 30.3<br>± 3.5 |
| 2.50 | 108 | 186.4<br>± 24.3 | 3.7<br>± 2.1 | 35.9<br>± 3.4 | 29 | 175.1<br>± 15.1 | 1.7<br>± 0.6 | 34.9<br>± 3.3 | 51 | 178.9<br>± 17.2 | 1.3<br>± 0.4 | 34.8<br>± 2.2 | 24 | 172.9<br>± 13.0 | 0.7<br>± 0.2 | 34.8<br>± 3.7 |
| 3.00 | 91 | 220.2<br>± 24.5 | 5.6<br>± 2.4 | 41.8<br>± 3.6 | 29 | 209.5<br>± 20.7 | 2.8<br>± 1.3 | 40.9<br>± 3.9 | 51 | 213.7<br>± 20.9 | 1.9<br>± 0.8 | 41.0<br>± 2.8 | 24 | 207.4<br>± 15.6 | 0.8<br>± 0.2 | 41.2<br>± 4.0 |
| 3.50 | 57 | 251.1<br>± 24.5 | 8.1<br>± 2.0 | 47.4<br>± 4.3 | 28 | 243.2<br>± 25.2 | 5.2<br>± 1.9 | 46.6<br>± 4.7 | 51 | 251.0<br>± 24.7 | 3.7<br>± 1.7 | 47.3<br>± 3.4 | 24 | 242.0<br>± 18.2 | 1.0<br>± 0.4 | 46.8<br>± 3.4 |
| 4.00 |  |  |  |  | 28 | 276.6<br>± 31.6 | 8.4<br>± 2.4 | 50.6<br>± 5.4 | 50 | 287.2<br>± 31.0 | 6.5<br>± 2.6 | 53.40<br>± 3.9 | 24 | 276.6<br>± 20.9 | 1.4<br>± 0.6 | 53.3<br>± 2.9 |
| 4.50 |  |  |  |  | 6 | 282.1<br>± 44.0 | 8.5<br>± 1.7 | 59.10<br>± 3.3 | 18 | 311.3<br>± 24.9 | 7.6<br>± 2.1 | 58.6<br>± 2.4 | 24 | 311.1<br>± 23.5 | 2.8<br>± 1.4 | 59.1<br>± 3.0 |
| 5.00 |  |  |  |  |  |  |  |  | 4 | 351.0<br>± 24.0 | 8.7<br>± 2.9 | 63.5<br>± 1.5 | 24 | 345.7<br>± 26.0 | 5.3<br>± 2.3 | 64.9<br>± 3.1 |
| 5.50 |  |  |  |  |  |  |  |  |  |  |  |  | 19 | 379.0<br>± 29.1 | 7.7<br>± 1.9 | 67.7<br>± 3.0 |
| 6.00 |  |  |  |  |  |  |  |  |  |  |  |  | 3 | 392.6<br>± 33.0 | 8.4<br>± 1.9 | 70.8<br>± 6.6 |

**Figure 1.**
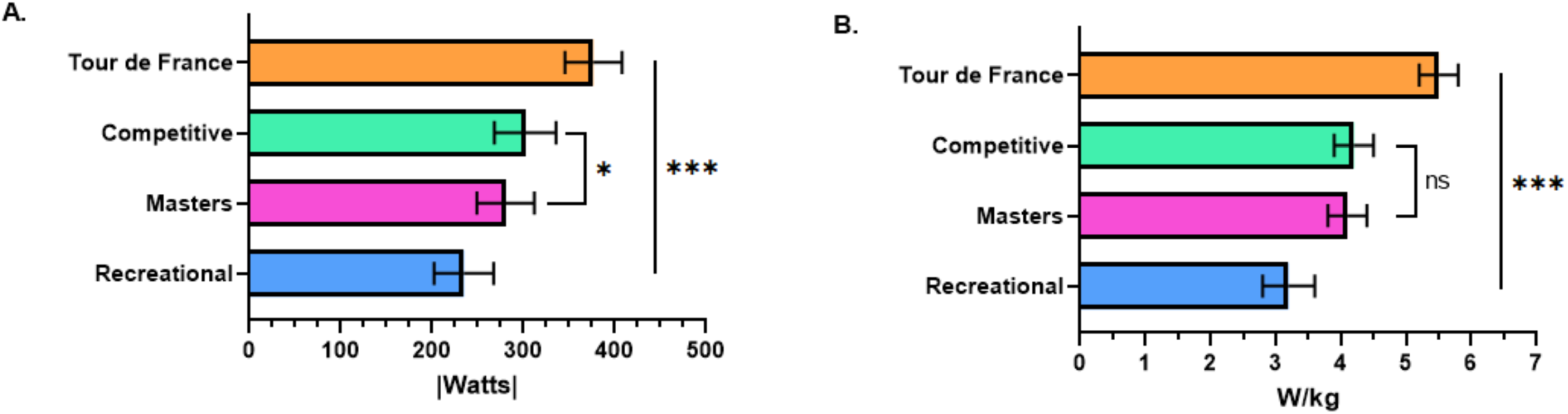
(A) Maximum absolute power output and (B) relative power output of each fitness category.

### 3.1 Cardiorespiratory parameters

Both absolute and relative VO2_max_ were also significantly higher in the TdF group than the rest of groups (p<0.001, Table 2-Figs. 2A-B). Likewise, VO₂ measured at 50% and 75% of VO₂max followed a similar hierarchical pattern, with TdF cyclists exhibiting the highest values at matched relative intensities (p<0.001), (Fig. 2C–D). Similar hierarchical patterns were found regarding total power output (W) relative to 50%, 70% and 100% of VO2_max_ (p<0.001) (Fig-2E). Both absolute and relative VO2_max_ highly correlated with absolute and relative maximal power respectively for all groups together as well as for each individual group (Figs 2F-G).

**Figure 2.**
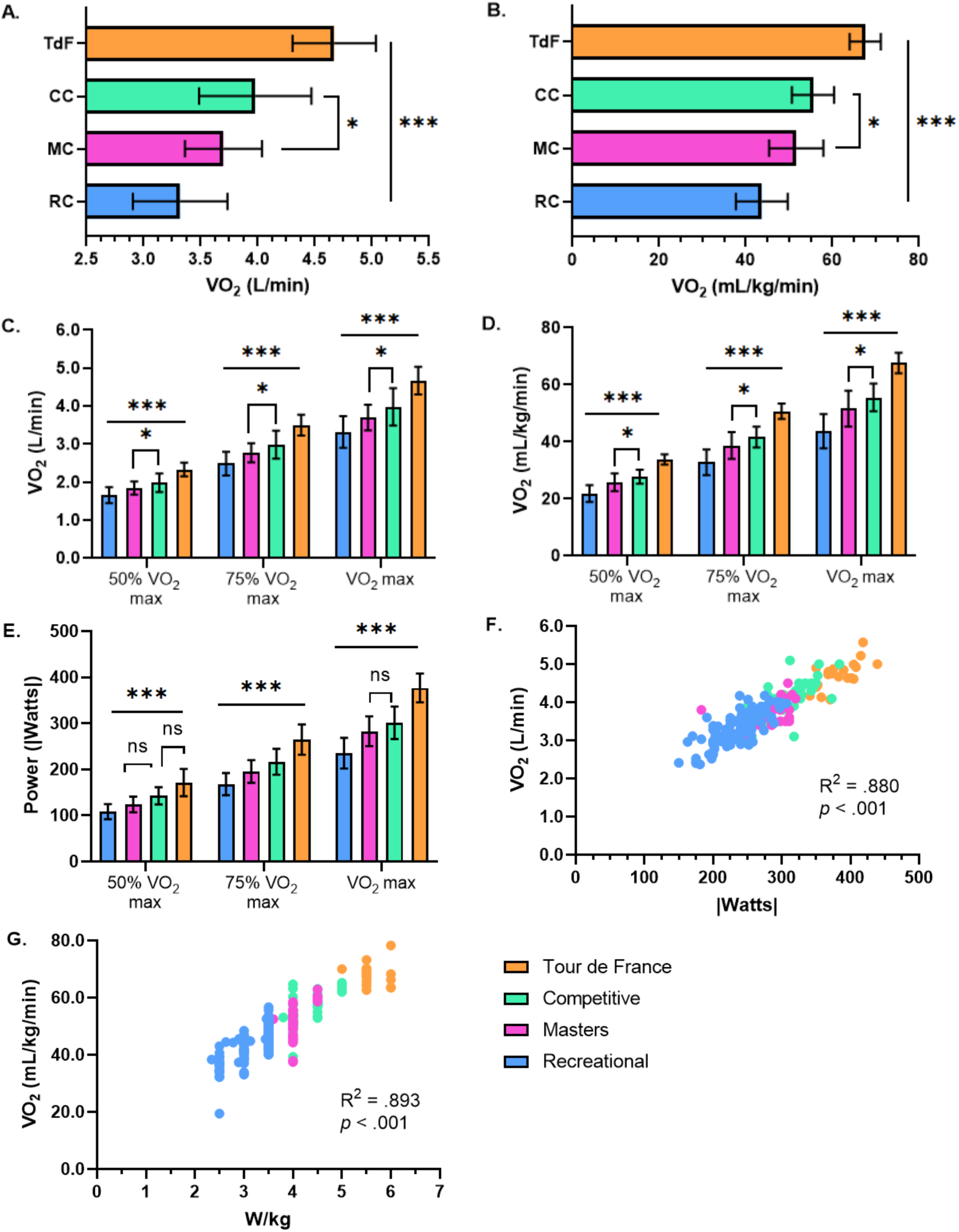
(A) Absolute and (B) relative VO2max for all fitness groups as well as (C) absolute and (D) relative VO2 at 50% VO2max, 75% VO2max, and 100% VO2max. (E) Absolute power output at 50%, 70 %, and 100% of VO2max. (F) Correlation between absolute VO2max and absolute max power output. (G) Correlation between relative VO2max and relative maximum power output for all fitness categories.

### 3.2 Metabolic parameters

#### 3.2.1. Blood lactate concentration

Blood lactate concentration was significantly lower across exercise intensities in TdF cyclists compared with all other groups at both relative and absolute power outputs (p < 0.001; Fig. 3A–B). Competitive cyclists also exhibited lower lactate concentrations than recreational cyclists, whereas differences between competitive and master cyclists were not statistically significant.

**Figure 3.**
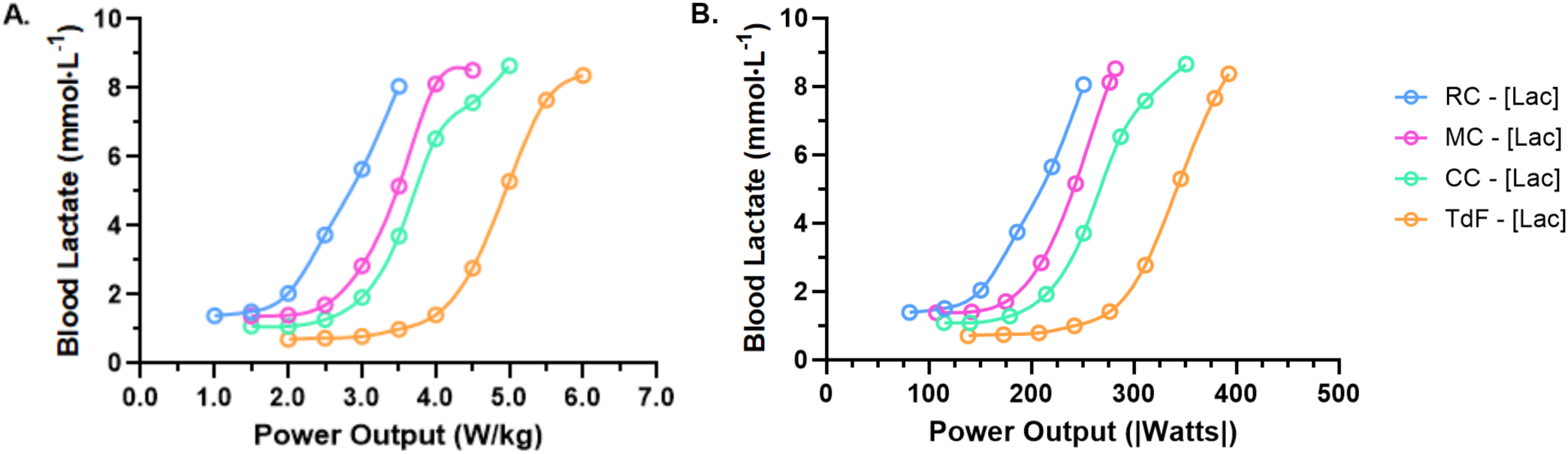
Mean blood lactate concentrations for both (A) relative power output and (B) absolute power output for the four groups of cyclists observed.

#### 3.2.2. Substrate Utilization

Fat oxidation rates were significantly higher in TdF cyclists across exercise intensities compared with all other groups at both relative and absolute power outputs (Fig. 4A–B).

**Figure 4.**
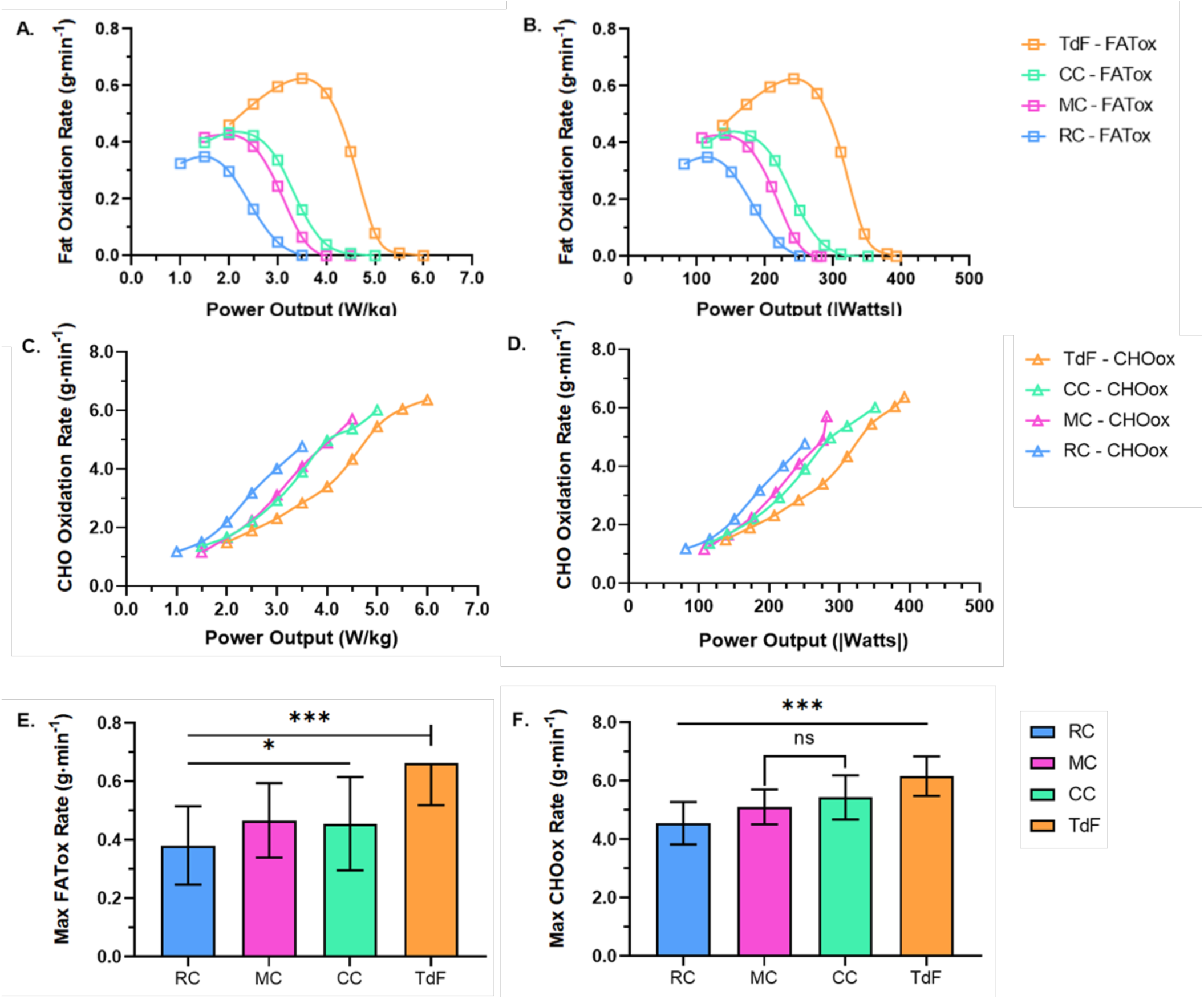
Mean fat oxidation rate in relation to increasing (A) relative power output and (B) absolute power output for the four groups studied. Mean carbohydrate oxidation rate in relation to increasing (C) relative power output and (D) absolute power output for the four groups studied. Comparison between (E) maximal fat oxidation rate and (F) maximal carbohydrate oxidation rate for the four groups studied.

Competitive cyclists demonstrated higher fat oxidation rates than master cyclists beginning at 3 W·kg⁻¹. At matched submaximal workloads, carbohydrate oxidation rates were lower in TdF cyclists compared with other groups (Fig. 4C–D).

At maximal exercise intensities, carbohydrate oxidation rates were highest in TdF cyclists, reflecting greater maximal metabolic demand. Both maximal fat oxidation (FATmax) and maximal carbohydrate oxidation (CHOmax) were significantly higher in TdF cyclists compared with all other groups (p < 0.001; Fig. 4E–F).

### 3.3. Correlations

Strong inverse correlations were observed between fat and carbohydrate oxidation rates across exercise intensities within each group and across all groups combined.

Similarly, blood lactate concentration exhibited strong inverse correlations with fat oxidation and strong positive correlations with carbohydrate oxidation across workloads.

These associations are descriptive and do not imply causality.

#### 3.3.1 CHOox vs FATox

We observed strong correlations between the average CHOox with the average FATox throughout the incremental test in all groups studied: TdF, r=−0.90, p<0.001; CC, r=−0.98, p=0.001; MC, r=−0.97, p<0.001; and RC, r=−0.98, p<0.001 (Fig-5). We also observed strong correlations between CHOox and FATox for all data points across the test within each group studied: TdF r=−0.77, p<0.001; CC, r=−0.82, p=0.001; MC, r=−0.86, p<0.001; and RC, r=−0.81, p<0.001 as well as for all data points of all groups together (r=−0.71, p<0.001) (Fig-6).

**Figure 5.**
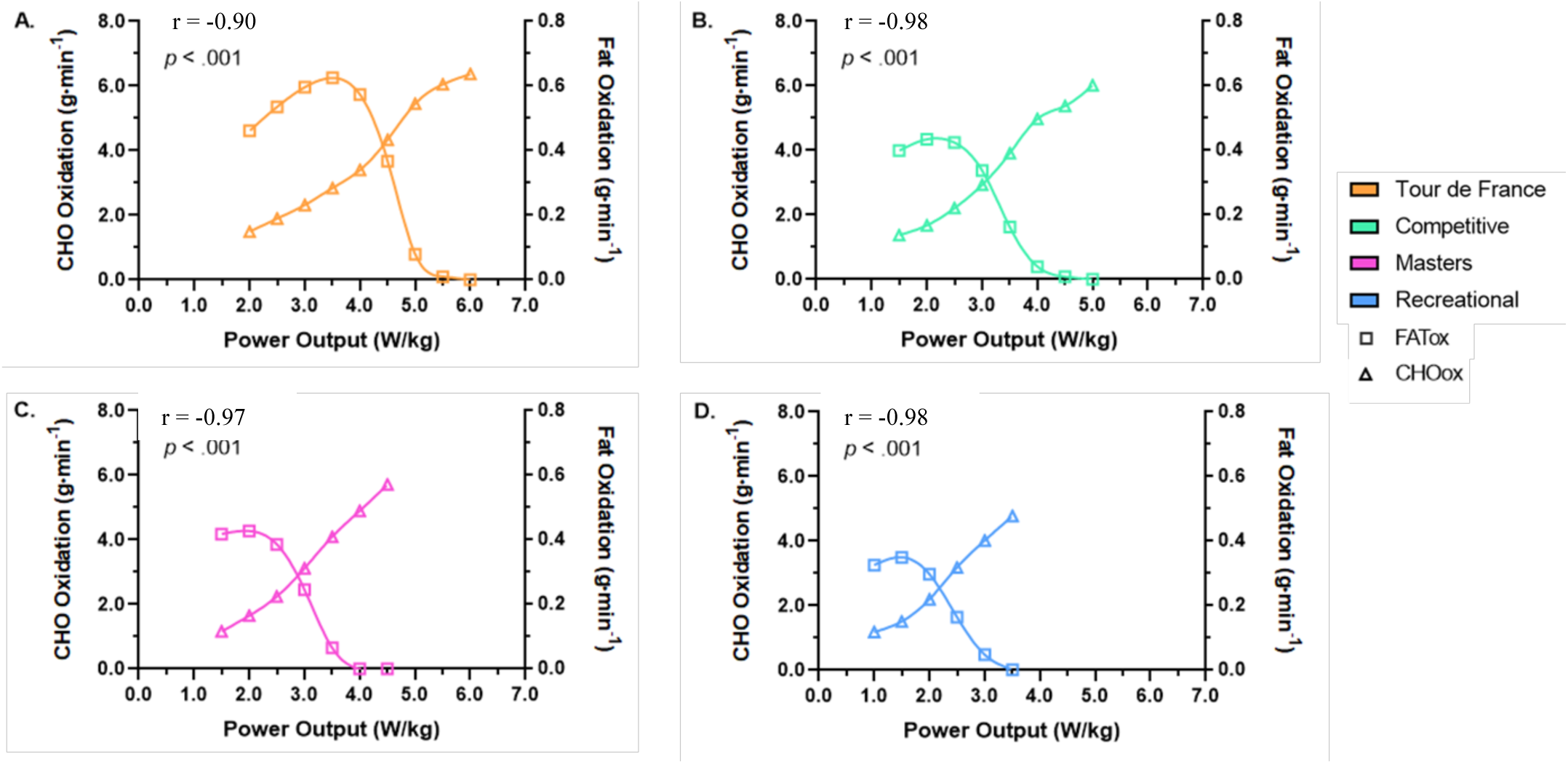
Mean carbohydrate oxidation rate and mean fat oxidation rate in relation to increasing workload in (A) Tour de France cyclists, (B) competitive cyclists, (C) competitive masters cyclists, and (D) recreational cyclists.

**Figure 6.**
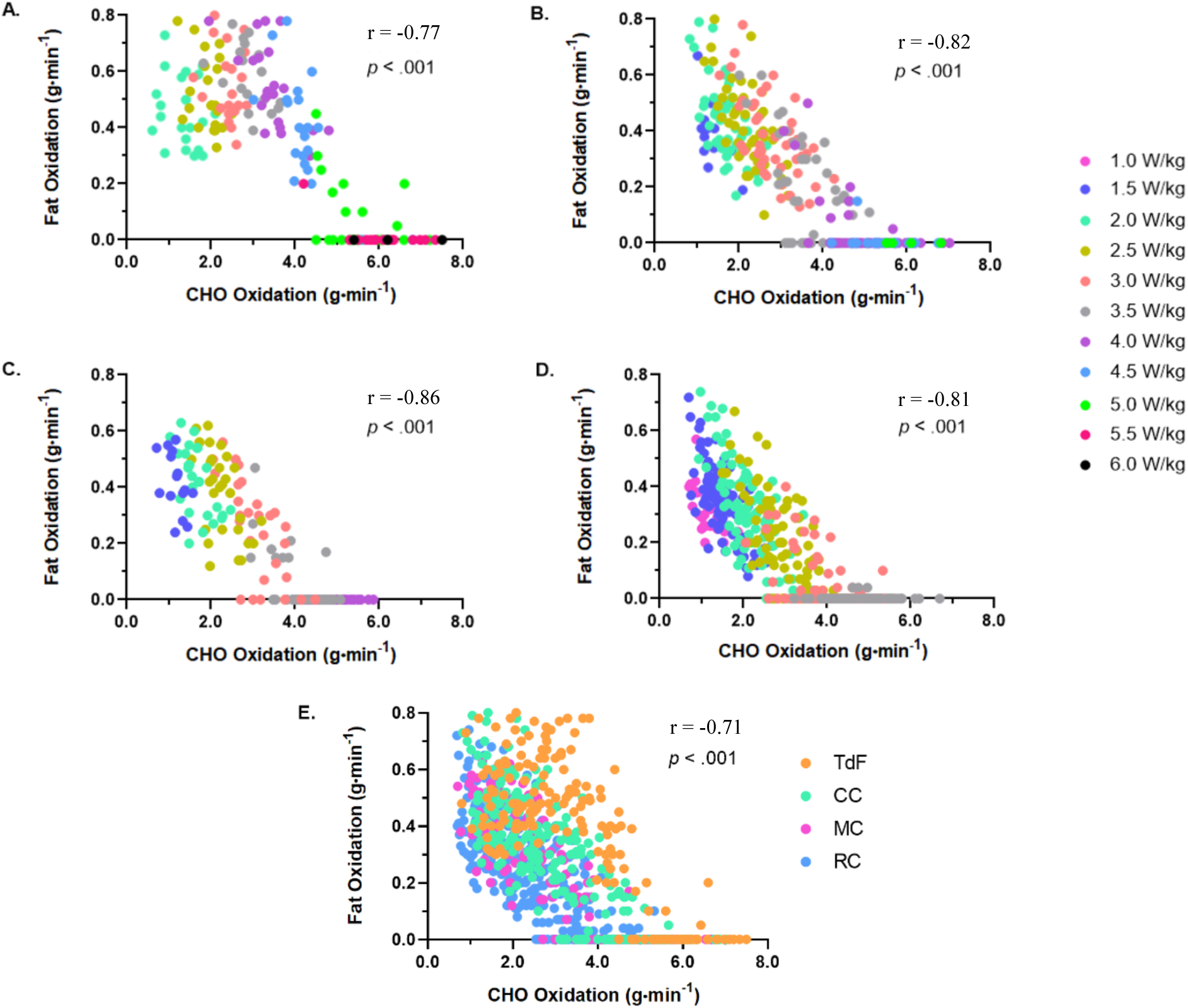
Correlations between fat oxidation rate and carbohydrate oxidation rate across all workloads in (A) Tour de France cyclists, (B) competitive cyclists, (C) competitive masters cyclists, and (D) recreational cyclists. (E) Correlation between fat oxidation rate and carbohydrate oxidation rate across all workloads for all four fitness groups.

#### 3.3.2. Blood Lactate vs FATox

The average correlations for [La^−^] with the average of FATox throughout the entire incremental test were especially strong in all groups studied: TdF, r=−0.96, p<0.001; CC, r=−0.97, p=0.001; MC, r=−0.96, p<0.001; and RC, r=−0.97, p<0.001 (Fig-7). We also observed strong correlations between [La^−^] and FATox for all data points across the test within each group studied: TdF r=−0.81, p<0.001; CC, r=−0.76, p=0.001; MC, r=−0.79, p<0.001; and RC, r=−0.76, p<0.001 as well as and for all data points of all groups together (r=−0.76, p<0.001) (Fig-8).

**Figure 7.**
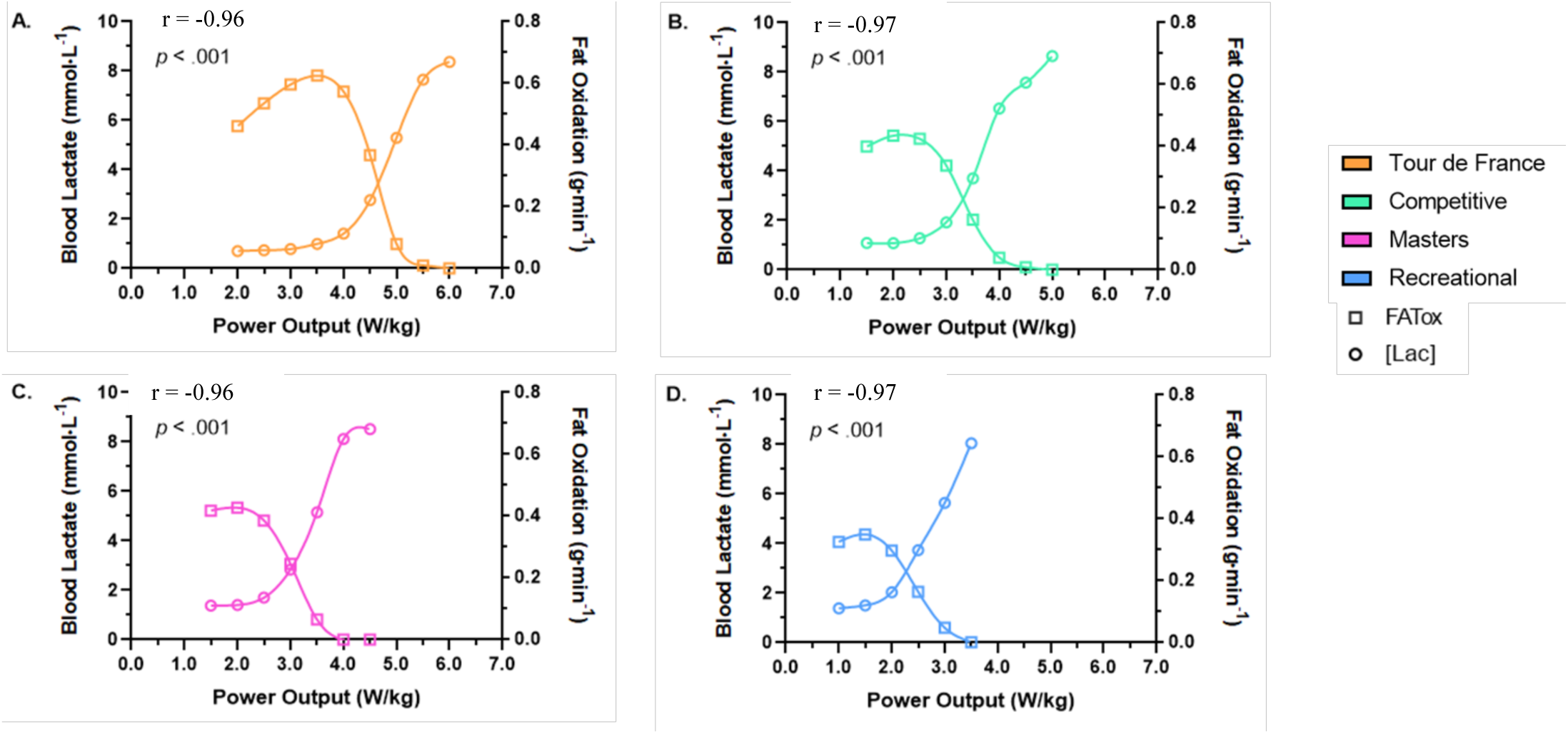
Mean blood lactate concentration and mean fat oxidation rate in relation to increasing workload in (A) Tour de France cyclists, (B) competitive cyclists, (C) competitive masters cyclists, and (D) recreational cyclists.

**Figure 8.**
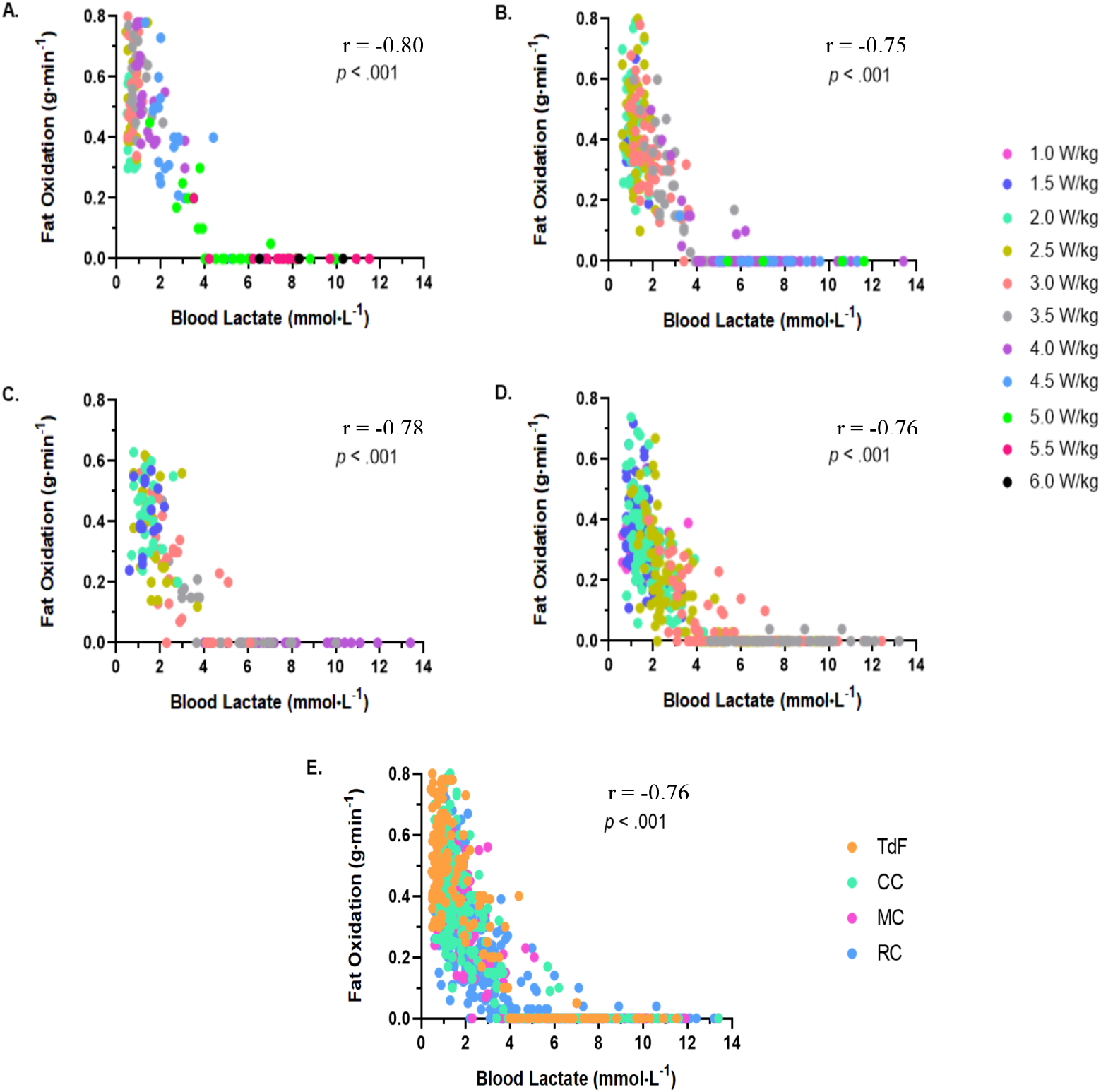
Correlations between fat oxidation rate and blood lactate concentration across all workloads in (A) Tour de France cyclists, (B) competitive cyclists, (C) competitive master cyclists, and (D) recreational cyclists. (E) Correlation between fat oxidation rate and blood lactate concentration across all workloads for all four groups studied.

#### 3.3.3. Blood Lactate vs CHOox

Finally, the correlations between average of [La^−^] and CHOox throughout the entire incremental test were also exceedingly strong in all groups; TdF, r=0.95, p<0.001; CC, r=0.98, p=0.001; MC, r=0.97, p<0.001; and RC, r=0.97, p<0.001 (Fig-9). We also observed strong correlations between [La^−^] and CHOox for all data points across the test within each group studied: TdF, r=0.87, p<0.001; CC, r=0.82, p=0.001; MC, r=0.84, p<0.001; RC, r=0.84, p<0.001; and r=0.80, p<0.001 for all data points of all groups together (Fig-10).

**Figure 9.**
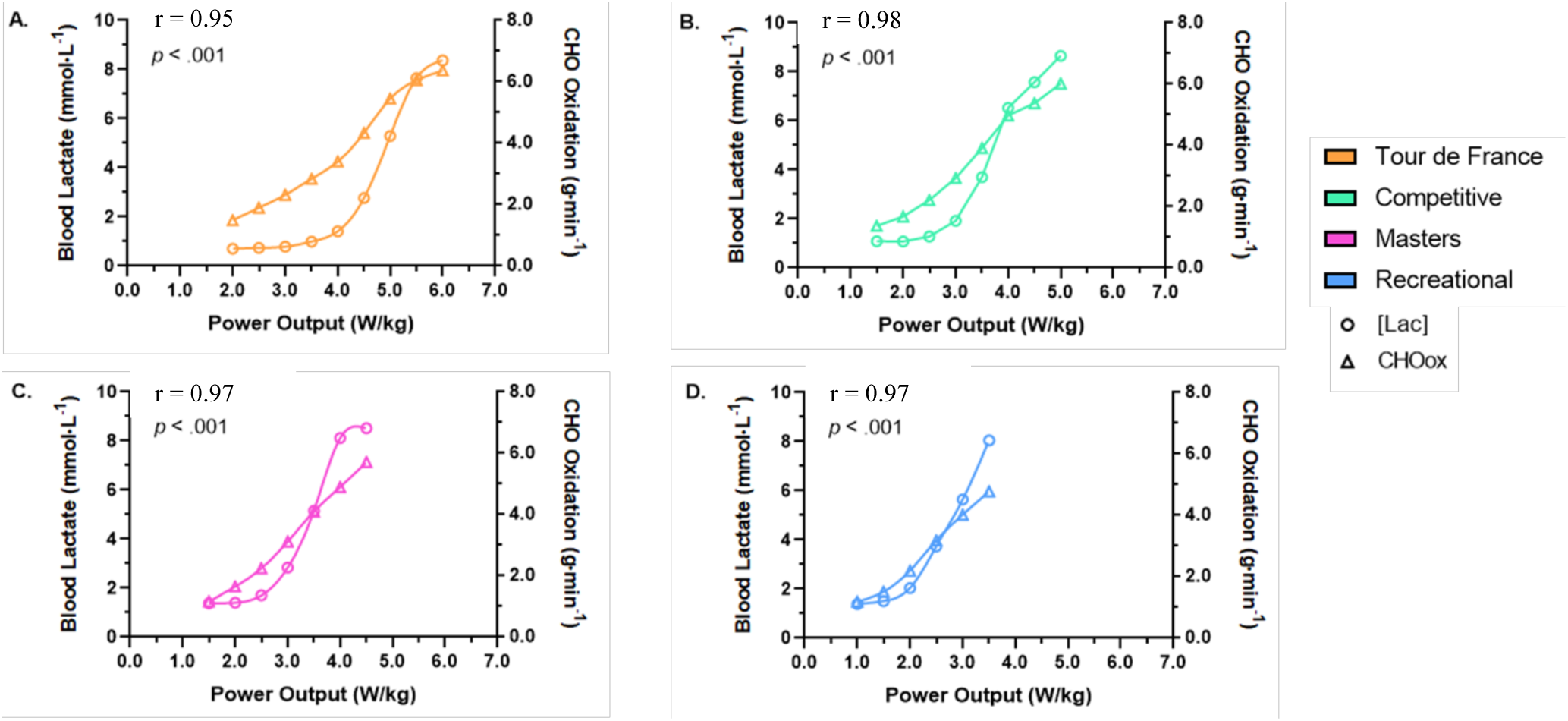
Mean blood lactate concentration and mean carbohydrate oxidation rate in relation to increasing workload in (A) Tour de France cyclists, (B) competitive cyclists, (C) competitive masters cyclists, and (D) recreational cyclists.

**Figure 10.**
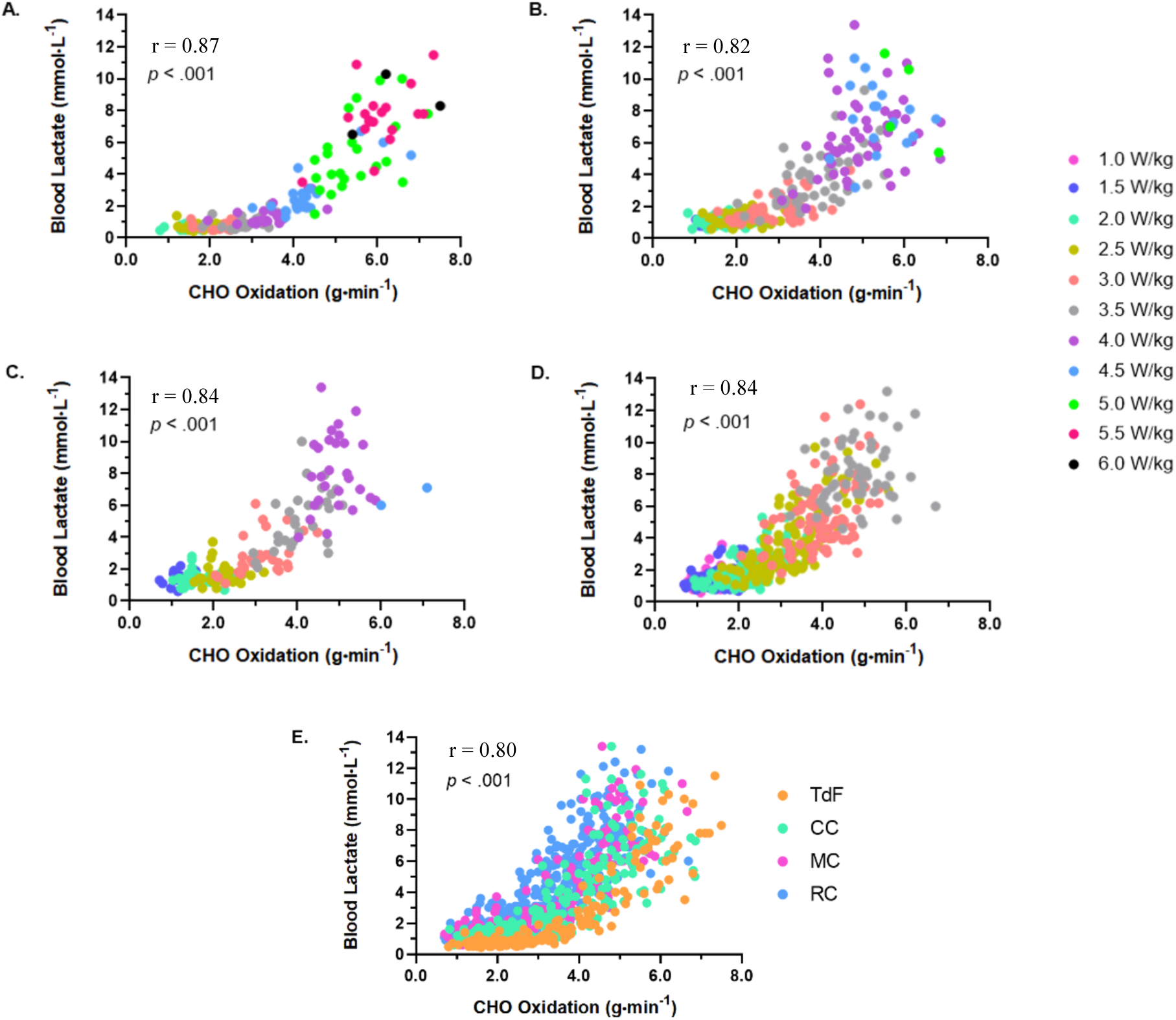
Correlations between carbohydrate oxidation rate and blood lactate concentration across all workloads in (A) Tour de France cyclists, (B) competitive cyclists, (C) competitive master cyclists, and (D) recreational cyclists. (E) Correlation between carbohydrate oxidation rate and blood lactate concentration across all workloads for all four groups studied.

## 4. DISCUSSION

Throughout this study, we demonstrate that Tour de France cyclists exhibit superior physiological responses to exercise compared with all other groups studied. Moreover, these physiological and metabolic responses follow a clear hierarchical pattern according to competitive level (TdF > CC ≥ MC > RC). While these findings are consistent with prior observations in elite endurance athletes, the present data provide a comprehensive comparison across a wide spectrum of fitness levels using integrated cardiorespiratory and metabolic measurements.

The assessment of cardiorespiratory fitness through measurement of maximal oxygen uptake (VO₂max) has been widely regarded as the gold standard for evaluating endurance capacity in both clinical and athletic settings. As anticipated, TdF cyclists demonstrated the highest VO₂max values, both in absolute and relative terms, and VO₂max correlated strongly with maximal power output across all groups. These findings reinforce the importance of VO₂max as a determinant of endurance performance while highlighting its close relationship with mechanical power production.

However, advances in exercise metabolism over recent decades have expanded the focus beyond cardiorespiratory capacity toward the metabolic and cellular determinants of performance. Consequently, the assessment of blood lactate concentration and substrate utilization during exercise has become increasingly common in applied physiology. In the present study, we characterized metabolic responses to exercise primarily through measurements of blood lactate concentration, fat oxidation, and carbohydrate oxidation across cyclists of varying fitness levels, in order to assess the metabolic responses to exercise which could lead to more developed exercise prescriptions than the information obtained from measuring VO2_max_ or even ventilatory thresholds historically provided.

In order to offer an updated perspective of the novel approaches to assess metabolic performance in athletes, the purpose of this study was to describe these metabolic responses to exercise in cyclists of different fitness levels, mainly via [La^−^] as well as by FATox and CHOox.

Lactate is the mandatory byproduct of glycolysis and probably the preferred fuel for most cells in the body, including skeletal muscle(Brooks, 2020). Thus, a high glycolytic rate, like in the case of high exercise intensities, always implies lactate production from glycolytic (fast-twitch) muscle fibers. The preferred route of lactate is to be shuttled into mitochondria in oxidative (slow-twitch) muscle fibers via cell-to-cell lactate shuttle (Brooks, 2009, Brooks, 2018) where lactate becomes a main fuel for oxidative phosphorylation in mitochondria(Brooks, 2018). When the rate of lactate production exceeds the capacity for oxidation, lactate accumulates within muscle and is released into the circulation. Accordingly, blood lactate concentration reflects the balance between glycolytic production and oxidative clearance during exercise. In this context, blood lactate concentration provides a practical, indirect reflection of integrated metabolic and bioenergetic responses during exercise including mitochondrial capacity to oxidize lactate(San-Millan and Brooks, 2018).

In parallel, fatty acids are oxidized within skeletal muscle mitochondria and contribute substantially to ATP production during submaximal exercise(Rasmussen and Wolfe, 1999). Thus, the indirect measurement of fat oxidation through stoichiometric equations based upon gas exchange (Frayn, 1983) could also represent a useful surrogate for the assessment of mitochondrial function(San-Millan and Brooks, 2018). Prior work has demonstrated regulatory interactions between lactate and lipid metabolism, including lactate-mediated inhibition of lipolysis via activation of GPR81 in adipose tissue(Liu et al., 2009). Further, as we have recently shown, lactate also elicits autocrine effects on fat metabolism by decreasing the activity in mitochondria of carnitine palmitoyltransferase I and II (CPT I and CPT II) in cardiomyocytes(San-Millan et al., 2022), therefore reducing mitochondrial fatty acid transport. As expected, TdF cyclists have a superior lactate clearance capacity and fat oxidation capacity compared to the rest of the groups studied, implying significantly more efficient mitochondrial function and bioenergetics.

At higher exercise intensities, carbohydrate oxidation increased in all groups. While TdF cyclists relied less on carbohydrate oxidation at matched submaximal workloads, they demonstrated the highest carbohydrate oxidation rates at maximal intensities, reflecting greater absolute metabolic demand and glycolytic capacity.

A notable observation of this study is the strong inverse relationship between blood lactate concentration and fat oxidation, along with the strong positive relationship between blood lactate concentration and carbohydrate oxidation across all fitness levels.

These associations were consistent across exercise intensities and competitive categories, underscoring the close coupling between lactate dynamics and substrate utilization during exercise. Importantly, these relationships are descriptive in nature and do not imply causality.

From an applied perspective, these findings have practical implications for physiological assessment and training prescription. While VO₂max testing provides valuable information, it requires specialized equipment and trained personnel, limiting accessibility. In contrast, blood lactate testing is relatively inexpensive, portable, and readily applicable in both laboratory and field settings. The strong associations observed between blood lactate concentration, substrate utilization, and performance-related variables suggest that lactate testing can provide meaningful insight into metabolic responses to exercise across a wide range of athletic populations.

When interpreted alongside workload and training context, lactate measurements may aid in characterizing metabolic efficiency and informing individualized exercise prescription.

In summary, this study provides a comprehensive characterization of physiological and metabolic responses to exercise across cyclists spanning a broad range of fitness levels, from Tour de France professionals to recreational athletes. Across all measured parameters, performance and metabolic responses followed a clear hierarchical pattern corresponding to competitive level.

While maximal oxygen uptake and power output remain important indicators of endurance performance, the present findings demonstrate that metabolic parameters provide additional and highly relevant information regarding exercise responses. Blood lactate concentration, fat oxidation, and carbohydrate oxidation exhibited strong and consistent relationships across exercise intensities and fitness categories.

Blood lactate concentration was closely associated with substrate utilization, demonstrating strong inverse relationships with fat oxidation and strong positive relationships with carbohydrate oxidation. These relationships highlight lactate as a practical, integrative indicator of metabolic responses to exercise.

From an applied perspective, the strong coupling between blood lactate concentration, substrate utilization, and workload supports the use of lactate-based testing to inform individualized exercise prescription. When interpreted alongside power output and training context, lactate measurements can assist in defining training intensities, monitoring metabolic adaptations, and optimizing training strategies across a wide range of athletic populations.

Given its affordability, portability, and applicability in both laboratory and field settings, lactate testing represents a valuable tool for both physiological assessment and exercise prescription, complementing traditional cardiorespiratory measurements and supporting more individualized and metabolically informed training programs.

## References

Achten, J. & Jeukendrup, A. E. 2003. Maximal fat oxidation during exercise in trained men. Int J Sports Med, 24, 603–8.

Achten, J., Venables, M. C. & Jeukendrup, A. E. 2003. Fat oxidation rates are higher during running compared with cycling over a wide range of intensities. Metabolism, 52, 747–52.

Benedict, F. G. 1938. Vital energetics. A study in comparative basal metabolism.

Bergman, B. C., Wolfel, E. E., Butterfield, G. E., Lopaschuk, G. D., Casazza, G. A., Horning, M. A. & Brooks, G. A. 1999. Active muscle and whole body lactate kinetics after endurance training in men. Journal of applied physiology (Bethesda, Md : 1985), 87, 1684–96.

Billat, L. V. 1996. Use of blood lactate measurements for prediction of exercise performance and for control of training. Recommendations for long-distance running. Sports Med, 22, 157–75.

Bonen, A., Mccullagh, K. J., Putman, C. T., Hultman, E., Jones, N. L. & Heigenhauser, G. J. 1998. Short-term training increases human muscle MCT1 and femoral venous lactate in relation to muscle lactate. Am J Physiol, 274, E102–7.

Brooks, G. A. 2009. Cell-cell and intracellular lactate shuttles. J Physiol, 587, 5591–600.

Brooks, G. A. 2018. The Science and Translation of Lactate Shuttle Theory. Cell metabolism, 27, 757–785.

Brooks, G. A. 2020. The tortuous path of lactate shuttle discovery: From cinders and boards to the lab and ICU. Journal of Sport and Health Science.

Donovan, C. M. & Brooks, G. A. 1983. Endurance training affects lactate clearance, not lactate production. The American journal of physiology, 244, E83–92.

Dubouchaud, H., Butterfield, G. E., Wolfel, E. E., Bergman, B. C. & Brooks, G. A. 2000. Endurance training, expression, and physiology of LDH, MCT1, and MCT4 in human skeletal muscle. American journal of physiology Endocrinology and metabolism, 278, E571–9.

Faulkner, J. A. 1966. Physiology of swimming. Res Q, 37, 41–54.

Fletcher, W. M. 1907. Lactic acid in amphibian muscle. J Physiol, 35, 247–309.

Frandsen, J., Vest, S. D., Larsen, S., Dela, F. & Helge, J. W. 2017. Maximal Fat Oxidation is Related to Performance in an Ironman Triathlon. Int J Sports Med, 38, 975–982.

Frayn, K. N. 1983. Calculation of substrate oxidation rates in vivo from gaseous exchange. J Appl Physiol Respir Environ Exerc Physiol, 55, 628–34.

Gollnick, P. D., Bayly, W. M. & Hodgson, D. R. 1986. Exercise intensity, training, diet, and lactate concentration in muscle and blood. Med Sci Sports Exerc, 18, 334–40.

Hill, A. V. A. L., H. 1923. Muscular exercise, lactic acid, and the supply and utilization of oxygen. An International Journal of Medicine, 62, 135–171.

Hoogeveen, A. R. 2000. The effect of endurance training on the ventilatory response to exercise in elite cyclists. Eur J Appl Physiol, 82, 45–51.

Impellizzeri, F. M., Ebert, T., Sassi, A., Menaspa, P., Rampinini, E. & Martin, D. T. 2008. Level ground and uphill cycling ability in elite female mountain bikers and road cyclists. Eur J Appl Physiol, 102, 335–41.

Jacobs, I., Sjodin, B. & Schele, R. 1983. A Single Blood Lactate Determination as an Indicator of Cycle Ergometer Endurance Capacity. European Journal of Applied Physiology and Occupational Physiology, 50, 355–364.

Knechtle, B., Muller, G., Willmann, F., Kotteck, K., Eser, P. & Knecht, H. 2004. Fat oxidation in men and women endurance athletes in running and cycling. Int J Sports Med, 25, 38–44.

Liu, C., Wu, J., Zhu, J., Kuei, C., Yu, J., Shelton, J., Sutton, S. W., Li, X., Yun, S. J., Mirzadegan, T., Mazur, C., Kamme, F. & Lovenberg, T. W. 2009. Lactate inhibits lipolysis in fat cells through activation of an orphan G-protein-coupled receptor, GPR81. J Biol Chem, 284, 2811–22.

Lucia, A., Hoyos, J., Pardo, J. & Chicharro, J. L. 2000a. Metabolic and neuromuscular adaptations to endurance training in professional cyclists: a longitudinal study. Jpn J Physiol, 50, 381–8.

Lucia, A., Joyos, H. & Chicharro, J. L. 2000b. Physiological response to professional road cycling: climbers vs. time trialists. Int J Sports Med, 21, 505–12.

Lucia, A., Pardo, J., Durantez, A., Hoyos, J. & Chicharro, J. L. 1998. Physiological differences between professional and elite road cyclists. Int J Sports Med, 19, 342–8.

Mcdermott, J. C. & Bonen, A. 1993. Endurance training increases skeletal muscle lactate transport. Acta physiologica Scandinavica, 147, 323–7.

O’toole, M. L., Douglas, P. S. & Hiller, W. D. 1989. Lactate, oxygen uptake, and cycling performance in triathletes. Int J Sports Med, 10, 413–8.

Padilla, S., Mujika, I., Cuesta, G. & Goiriena, J. J. 1999. Level ground and uphill cycling ability in professional road cycling. Med Sci Sports Exerc, 31, 878–85.

Padilla, S., Mujika, I., Orbananos, J., Santisteban, J., Angulo, F. & JOSE Goiriena, J. 2001. Exercise intensity and load during mass-start stage races in professional road cycling. Med Sci Sports Exerc, 33, 796–802.

Randell, R. K., Rollo, I., Roberts, T. J., Dalrymple, K. J., Jeukendrup, A. E. & Carter, J. M. 2017. Maximal Fat Oxidation Rates in an Athletic Population. Med Sci Sports Exerc, 49, 133–140.

Rasmussen, B. B. & Wolfe, R. R. 1999. Regulation of fatty acid oxidation in skeletal muscle. Annu Rev Nutr, 19, 463–84.

San-Millan, I. & Brooks, G. A. 2018. Assessment of Metabolic Flexibility by Means of Measuring Blood Lactate, Fat, and Carbohydrate Oxidation Responses to Exercise in Professional Endurance Athletes and Less-Fit Individuals. Sports Med, 48, 467–479.

San-Millan, I., Gonzalez-Haro, C. & Sagasti, M. 2009. Physiological Differences Between Road Cyclists of Different Categories: A New Approach. Med. Sci. Sports Exerc, 41, 64–65.

San-Millan, I., Sparagna, G. C., Chapman, H. L., Warkins, V. L., Chatfield, K. C., Shuff, S. R., Martinez, J. L. & Brooks, G. A. 2022. Chronic Lactate Exposure Decreases Mitochondrial Function by Inhibition of Fatty Acid Uptake and Cardiolipin Alterations in Neonatal Rat Cardiomyocytes. Front Nutr, 9, 809485.

Shephard, R. J., C. Allen, A. J. S. Benade, C. T. M. Davies, P. E. DI Prampero, R. Hedman, J. E. Merriman, K. Myhre, and R. Simmons 1968. The maximum oxygen intake: An international reference standard of cardio-respiratory fitness. Bulletin of the World Health Organization, 38, 757.

Stisen, A. B., Stougaard, O., Langfort, J., Helge, J. W., Sahlin, K. & Madsen, K. 2006. Maximal fat oxidation rates in endurance trained and untrained women. Eur J Appl Physiol, 98, 497–506.

Underwood, E. A. 1944. Lavoisier and the History of Respiration. Proc R Soc Med, 37, 247–62.

